# A New Suite of Allelic Exchange Vectors for the Scarless Modification of Proteobacterial Genomes

**DOI:** 10.1101/624551

**Authors:** Jacob E. Lazarus, Alyson R. Warr, Carole J. Kuehl, Rachel T. Giorgio, Brigid M. Davis, Matthew K. Waldor

**Affiliations:** Division of Infectious Digmseases, Massachusetts General Hospital, Boston, MA, USA; Department of Microbiology, Harvard Medical School, Boston, MA, USA; Division of Infectious Diseases, Brigham and Women’s Hospital, Boston, MA, USA; Howard Hughes Medical Institute, Boston, MA, USA

**Keywords:** Allelic exchange, toxin-antitoxin, Type VI toxin, *sacB*, *rhaS*, rhamnose induction, *amilCP*, *tsPurple*, *Serratia marcescens*, *Shigella flexneri*, *ampC*, *ampD*, *amiD*, peptidoglycan, amidohydrolase, beta-lactamase, antibiotic resistance

## Abstract

Despite the advent of new techniques for genetic engineering of bacteria, allelic exchange through homologous recombination remains an important tool for genetic analysis. Currently, *sacB*-based vector systems are often used for allelic exchange, but counter-selection escape, which prevents isolation of cells with the desired mutation, limits its utility. To circumvent this limitation, we engineered a series of “pTOX” allelic exchange vectors. Each plasmid encodes one of a set of inducible toxins, chosen for their potential utility in a wide range of medically important Proteobacteria. A codon-optimized *rhaS* transcriptional activator with a strong synthetic ribosome binding site enables tight toxin induction even in organisms lacking an endogenous rhamnose regulon. Expression of the blue *amilCP* or magenta *tsPurple* non-fluorescent chromoproteins facilitates monitoring of successful single- and double-crossover events using these vectors. The versatility of these vectors was demonstrated by deleting genes in *Serratia marcescens*, *Escherichia coli* O157:H7, *Enterobacter cloacae*, and *Shigella flexneri*. Finally, pTOX was used to characterize the impact of disruption of all combinations of the 3 orthologous *S. marcescens* peptidoglycan amidohydrolases on chromosomal *ampC* beta-lactamase activity and corresponding beta-lactam antibiotic resistance. Mutation of multiple amidohydrolases was necessary for high level *ampC* derepression and beta-lactam resistance. These data suggest why beta-lactam resistance may emerge during treatment less frequently in *S. marcescens* than in other AmpC-producing pathogens like *E. cloacae.* Collectively, our findings suggest that the pTOX vectors should be broadly useful for genetic engineering of Gram-negative bacteria.

**Importance:** Targeted modification of bacterial genomes is critical for genetic analyses of microorganisms. Allelic exchange is a technique that relies on homologous recombination to substitute native loci for engineered sequences. However, current allelic exchange vectors often enable only weak selection for successful homologous recombination. We developed a suite of new allelic exchange vectors, pTOX, which were validated in several medically important Proteobacteria. They encode visible non-fluorescent chromoproteins that enable easy identification of colonies bearing integrated vector, and permit stringent selection for the second step of homologous recombination, yielding modified loci. We demonstrate the utility of these vectors by using them to investigate the effect of inactivation of *Serratia marcescens* peptidoglycan amidohydrolases on beta-lactam antibiotic resistance.

## Introduction

The ever-increasing availability of bacterial genome sequence data has driven the demand for widely applicable and facile techniques enabling site-specific targeted mutagenesis. In general, such techniques can be divided into those that rely on exogenous enzymes versus those that depend exclusively on endogenous enzymes. Examples of methods in the former category include those utilizing the Lambda Red recombinase (“recombineering” (1)), those employing clustered regularly interspaced short palindromic repeat (CRISPR)/Cas9 systems (2), or a combination of the two (3, 4). These systems can be fast and reliable, but often require organism-specific modifications, rely on efficient transformation, and can leave genetic scars or result in off-target mutations.

In contrast, “allelic exchange” utilizes endogenous homologous recombination enzymes to facilitate the replacement of a native genomic region with a foreign sequence of interest. This is a versatile technique that can routinely yield mutations ranging from kilobase-scale deletions or insertions to the generation of precise point mutations. The early allele exchange vectors resulted in antibiotic-marked strains (5, 6); subsequent advances using counter-selectable cassettes allowed the generation of truly scarless, unmarked mutant strains. However, many genes used in counter-selection strategies (*e.g. rpsL*, *pheS, thyA, ccdB)* require a specific host genotype, limiting their widespread utility (7). Background-independent counter-selection strategies utilizing *tetAR* (8), *sacB* (9), or a combination of the two (10) are valuable but often require considerable optimization. Moreover, counter-selection escape, where the integrated allelic exchange vector remains lodged in the genome, preventing isolation of the desired mutant, remains common with such schemes even after optimization. This has been a key technical obstacle limiting wider use of allelic exchange.

Recently, a powerful negative selection system using inducible toxins derived from toxin-antitoxin systems or from Type VI secreted effector-toxins was developed for use with recombineering (11). Here, we repurpose these toxins for use in allelic exchange and engineer a counter-selection escape surveillance system using visible chromoproteins derived from the *Acropora millepora* coral. We demonstrate the utility of these new allele exchange vectors, designated “pTOX,” in multiple medically important Proteobacteria. These vectors were used to systematically delete all combinations of the three peptidoglycan hydrolases in *Serratia marcescens* to characterize their contributions to beta-lactam antibiotic resistance.

## Results

### Engineering and testing of pTOX vectors

The motivation for this work arose from our difficulties adapting common genetic tools for use in *Serratia marcescens,* an Enterobacteria that is a common cause of healthcare-associated urinary tract infections, pneumonia, and bacteremia (12). While chemical- and electro-transformation is possible in many *S. marcescens* strains (13, 14), it is often cumbersome and inefficient, which reduces the utility of Lambda red recombinase- and CRISPR/Cas9-based systems for genetic manipulation. Because of this, we sought to construct a conjugatable allelic exchange vector for *S. marcescens* that would be widely useful.

Our set of new vectors (the pTOX vectors) is derived from pDS132, a *sacB*-based suicide plasmid that contains the conditional (π-dependent) R6K origin of replication (15). The *sacB* cassette was replaced with a rhamnose-inducible toxin obtained from the pSLC vector series (11) (Fig 1A). Reasoning that a given toxin would be most useful in a strain that did not encode a chromosomal copy of that same toxin (and presumably the corresponding antitoxin or immunity protein), we identified a minimal set of three toxins (*yhaV, mqsR,* and *tse2*), of which at least one should be effective in the majority of medically important proteobacteria (Fig S1). Additional steps in the construction of this set of vectors included: 1) Introduction of a codon-optimized *rhaS* transcriptional activator (16) with a strong synthetic ribosome binding site (17) to enable use of the well-characterized and stringent rhamnose-inducible system for toxin activation (18) even in strains that lack a rhamnose regulon; 2) introduction of a strong forward transcriptional terminator upstream of the multiple cloning site, minimizing read-through into the multiple cloning site and facilitating the manipulation of toxic genes; and 3) introduction of a greatly expanded polylinker region (19) (Fig 1A) to facilitate insertion of new sequences into the vectors. Two versions of this set of plasmids, encoding either chloramphenicol or gentamicin resistance, were created (Supp Table 2). All molecular cloning was performed in the presence of glucose, which inhibits toxin production through catabolite repression.

**Fig 1.**
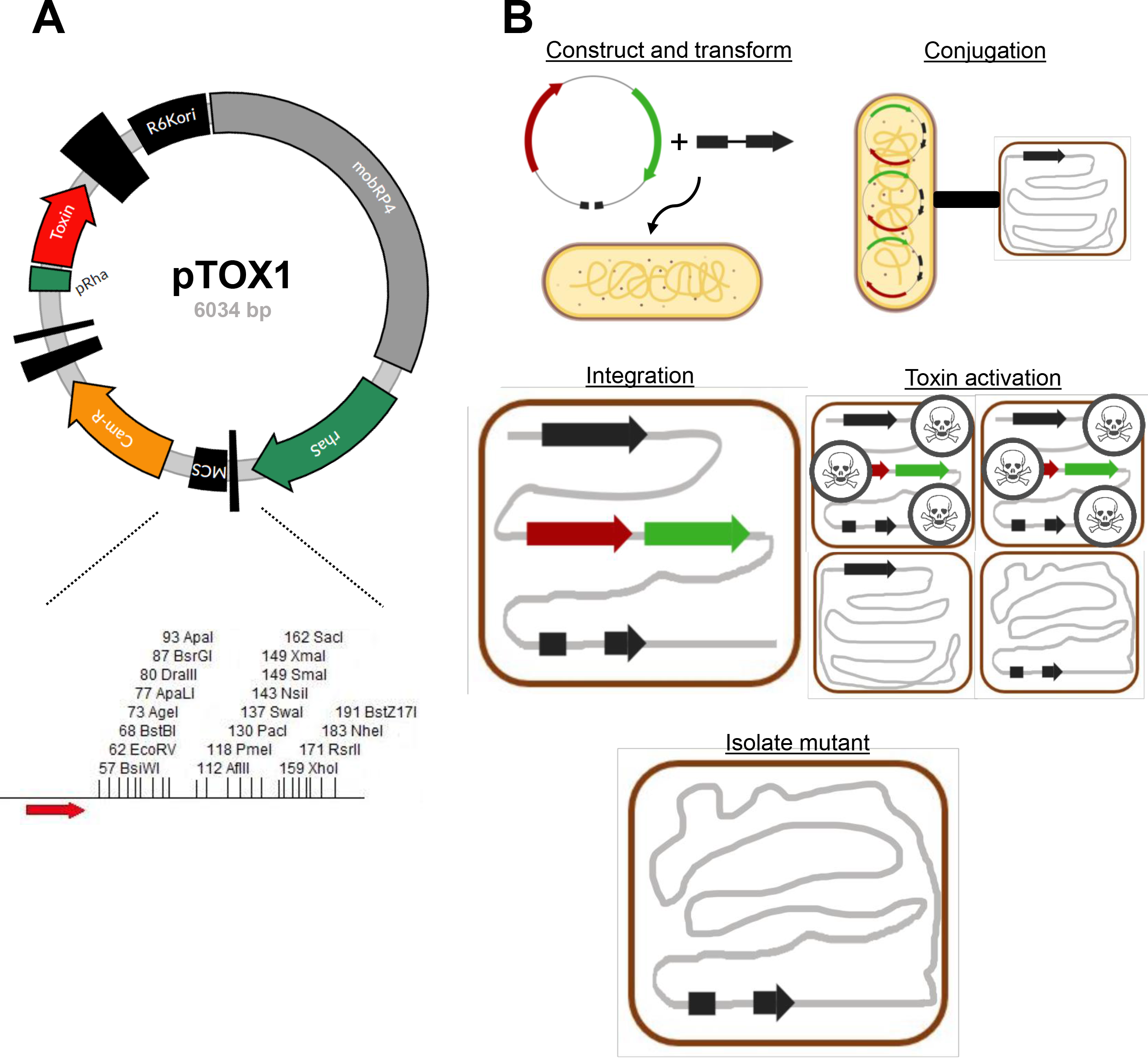
Allelic exchange with pTOX. A) Plasmid map of pTOX1. R6Kori, the R6K origin of replication; mobRP4, mobilization region from RP4 conjugative plasmid; *rhaS*, the rhamnose transcriptional activator; MCS, multiple cloning site; Cam-R, chloramphenicol resistance cassette; pRha, rhamnose promoter. Vertical black bars of varying width represent terminators. Bottom, expanded polylinker with restriction sites unique to pTOX1 (*yhaV*) shown. Red arrow, forward transcriptional terminator. B) pTOX workflow. Step 1: the desired allele is inserted into the MCS using isothermal assembly and transformed into donor *E. coli*. (yellow bacillus) Step 2: conjugation is performed between the donor *E. coli* and the organism of interest (red coccobacillus). Step 3: pTOX integrates into the appropriate chromosomal locus. Step 4: merodiploids are isolated and toxin induced. Step 5: the desired clone is identified by colony PCR.

The utility of each of the three toxins was validated in *S. marcescens* ATCC 13880, which lacks *rhaS* and endogenous versions of the 3 toxins. First, a region homologous to the targeted chromosomal locus was inserted into the pTOX multiple cloning site (see the Methods for more detail and Fig 1B for a schematic). Next, conjugation was used to introduce pTOX derivatives into *S. marcescens.* Single cross-over merodiploids were selected on the appropriate antibiotic. To assess the utility of the heterologous *rhaS*, we then compared the growth of merodiploids to wild-type *S. marcescens* in either glucose- or rhamnose-containing media. Toxin-containing merodiploids grown in glucose-containing media grew indistinguishably from wild-type, while growth in rhamnose-containing media was undetectable (Fig 2). These observations reveal that *yhaV*, *mqsR*, and *tse2* enable robust growth inhibition in *S. marcescens* and that the exogenous *rhaS* is sufficient for stringent control of their expression.

**Fig 2.**
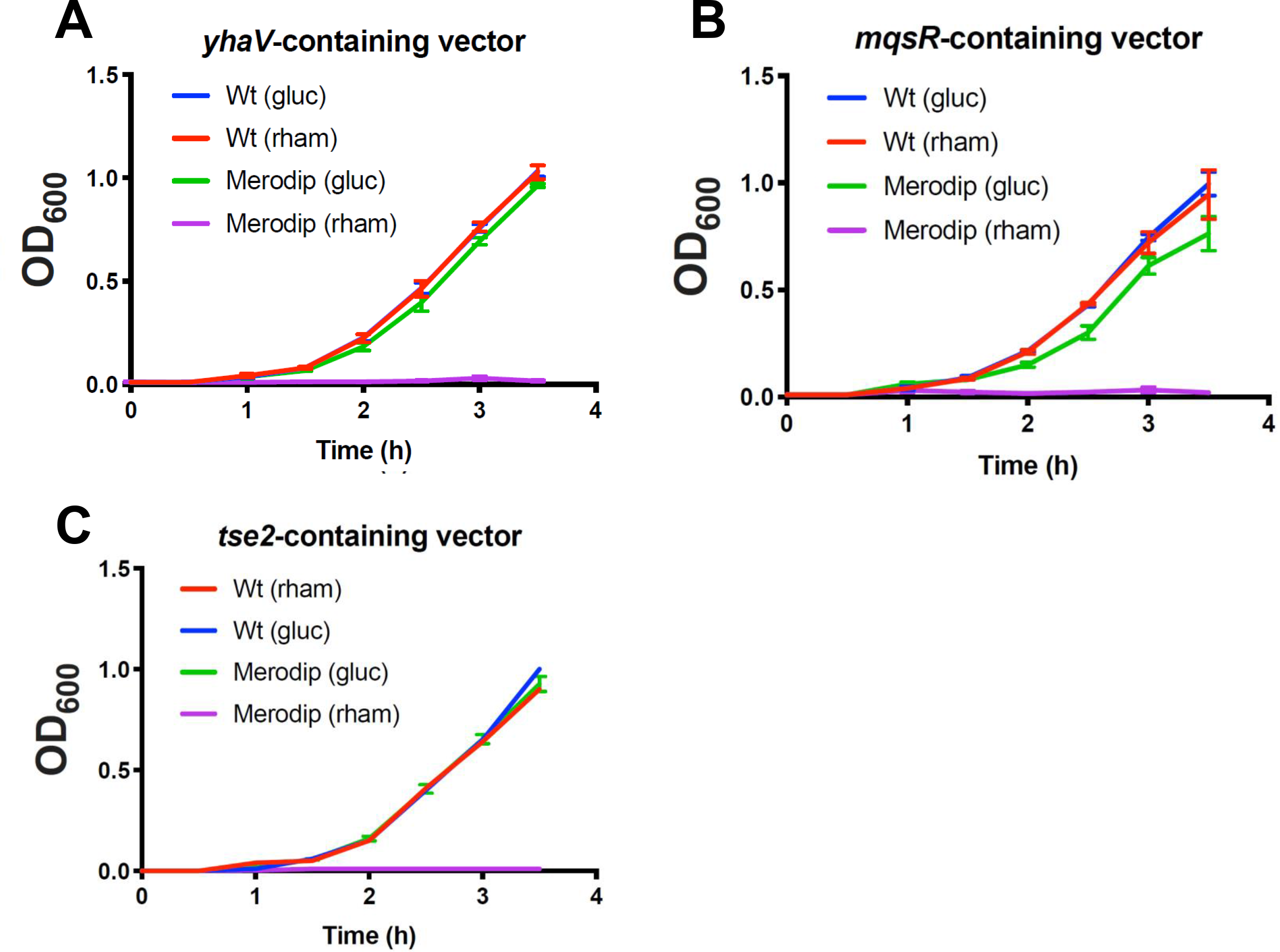
Induction of specific bacterial toxins inhibit *S. marcescens* growth. *S. marcescens* wild-type (Wt) or merodiploid (merodip) harboring the indicated pTOX-carrying toxin were diluted from exponential phase growth in LB into either 2% (w/v) glucose (gluc) or rhamnose-containing (rham) LB and incubated with agitation at 37°C. Note that the Wt (gluc) curve is obscured by the Wt (rham) curve in A and the error bars in C are smaller than the line for all but the merodiploid (gluc). Means and SEM are depicted from at least 3 independently generated merodiploids.

A limitation of the *sacB* counter-selection system is the occasional outgrowth of merodiploids that have either mutated the *sacB* gene or acquired resistance to its product (9). Such counter-selection escape can confound isolation of double-crossover events. To assess whether counter-selection escape also confounds *yhaV*-, *mqsR*-, or *tse2*-based selections, we randomly selected 23 colonies representing putative double-crossovers (based on growth in the presence of rhamnose) from 3 independent experiments for each of the three toxin-vectors. All 207 colonies screened were chloramphenicol sensitive and lacked pTOX vector sequences by PCR (Table 1 and Fig S2). These observations suggest that selection mediated by the 3 toxins is potent and that the frequency of counter selection escape is very low.

**TABLE 1.** Absence of integrated pTOX in putative double-crossover colonies^a^

|  | YhaV toxin | MqsR toxin | Tse2 toxin |
| --- | --- | --- | --- |
| **CAM<sup>R</sup>** | 0/69 | 0/69 | 0/69 |
| **PCR<sup>+</sup>** | 0/69 | 0/69 | 0/69 |

<sup>a</sup> CAM = chloramphenicol resistance was determined using the parental single-
crossover colony as a positive control. An intergenic region of pTOX was used for
colony PCR to avoid false negatives during selection for inactivated open reading
frames.

### Utility of pTOX vectors in diverse pathogens

To investigate the versatility of the pTOX vectors, we tested their capacity to mediate diverse allele replacements, beginning with the *S. marcescens hexS* locus. *S. marcescens* ATCC 13880, like many isolates of this opportunistic pathogen, produces the red prodigiosin pigment; however, production is only robust at reduced temperatures, due to relief of repression mediated by the negative regulator *hexS* (20). A pTOX derivative encoding sequences flanking *hexS* was used to delete this regulator from the *S. marcescens* chromosome, resulting in prodigiosin hyperproduction even at 37°C (Fig 3A). Subsequently, we have replaced more than 20 loci in *S. marcescens* using pTOX1, pTOX2, and pTOX3 (Supplemental Table 2). All attempts have been successful, though like other allelic exchange methods, the ratio of wild-type to mutant double-crossovers can vary from balanced to skewed.

**Fig 3.**
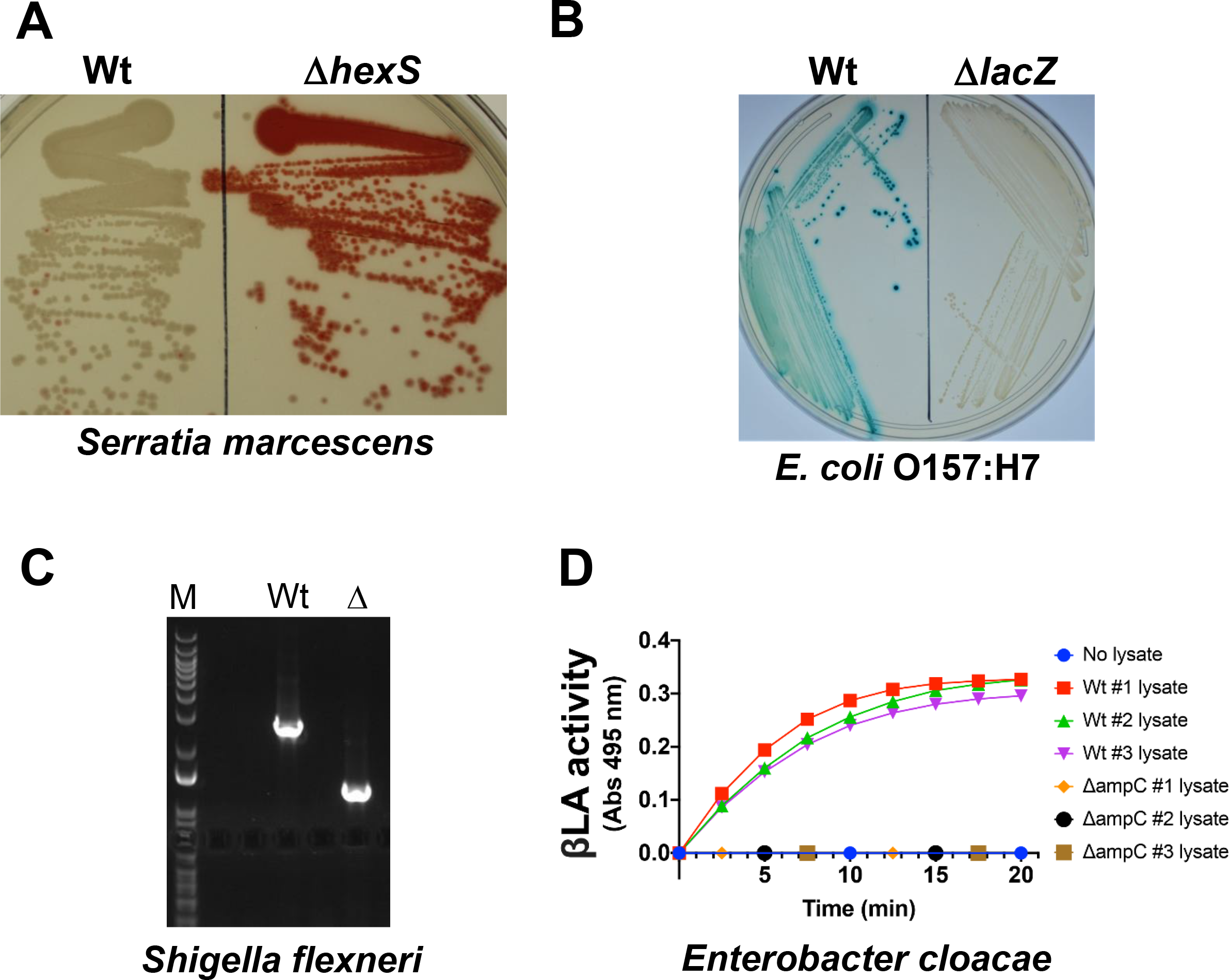
pTOX for genomic modification in multiple pathogens. A) *S. marcescens* colony coloration in Wt (left) *and* Δ*hexS* (right) grown at 37°C for 1 day. HexS inhibits expression of the red prodigiosin characteristic of *S. marcescens.* B) *E. coli* O157:H7 colony coloration in Wt (left) and Δ*lacZ* (right) grown on X-gal-containing media. Blue-green colony color indicates lactose fermentation. C) *S. flexneri* colony PCR and results of 1% agarose gel electrophoresis demonstrating deletion of *ipgH* from *S. flexneri* virulence plasmid. M, marker; Wt, wild-type; Δ, Δ*ipgH*. D) *E. cloacae* beta-lactamase activity in total clarified sonicate from 3 Wt double-crossover colonies and from 3 Δ*ampC* colonies. Sonicates were incubated with nitrocefin, a chromogenic cephalosporin substrate which absorbs at 495 nm when hydrolyzed.

We also tested the utility of the pTOX vectors in 3 additional Gram-negative pathogens. *Escherichia coli* O157:H7 (EHEC) is an important cause of foodborne diarrhea as well as a systemic microangiopathy which can lead to hemolysis and renal failure. A pTOX3 derivative was used to delete *lacZ,* which produces a beta-galactosidase that enables wild-type EHEC to ferment lactose. As seen in Fig 3B, deletion of EHEC *lacZ* yielded colonies that are white on agar containing the chromogenic lactose analog 5-bromo-4-chloro-3-indolyl-β-D-galactopyranoside (X-gal). Derivatives of pTOX3 were also used to replace nearly 20 additional loci in EHEC.

*Shigella flexneri* is an increasingly antibiotic-resistant cause of dysentery. In *S. flexneri*, most secreted virulence proteins (effectors) are encoded by a large, unstable virulence plasmid. Recombineering is useful in performing single gene deletions on the plasmid, but multiple gene deletions leave identical scar sequences that can enable undesired recombination within the plasmid. pTOX3 was used to delete the virulence plasmid *ipgH* locus (Fig 3C) as well as chromosomal loci.

Finally, pTOX was efficacious in *Enterobacter cloacae,* an opportunistic hospital-associated pathogen associated with urinary tract and bloodstream infections. *E. cloacae,* like *S. marcescens,* possesses an inducible chromosomal beta-lactamase, AmpC, which hydrolyzes most beta-lactam antibiotics. A pTOX3 derivative was used to delete *E. cloacae ampC*. Colonies harboring the *ampC* deletion exhibited no detectable beta-lactamase activity, whereas colonies that reverted to wild-type *(ampC^+^)* did (Figure 3D). Collectively, these observations suggest that pTOX may be widely useful in Gram-negative bacteria, particularly for those where other methods are difficult or unavailable.

### Chromoproteins facilitate visual detection of pTOX transconjugants

Conjugation efficiency can vary between species and strains. For organisms like *S. marcescens,* in which conjugation can be inefficient, we incorporated an additional module coding for the AmilCP protein into the pTOX vectors (Fig 4A). AmilCP is a non-fluorescent blue chromoprotein derived from the *Acropora millepora* coral; we sought to use its blue coloration as an additional method to discriminate wild-type colonies from transconjugants. To this aim, multiple combinations of promoters and ribosomal binding sites were tested to identify those which provided coloration sufficient for discrimination without special equipment.

**Fig 4.**
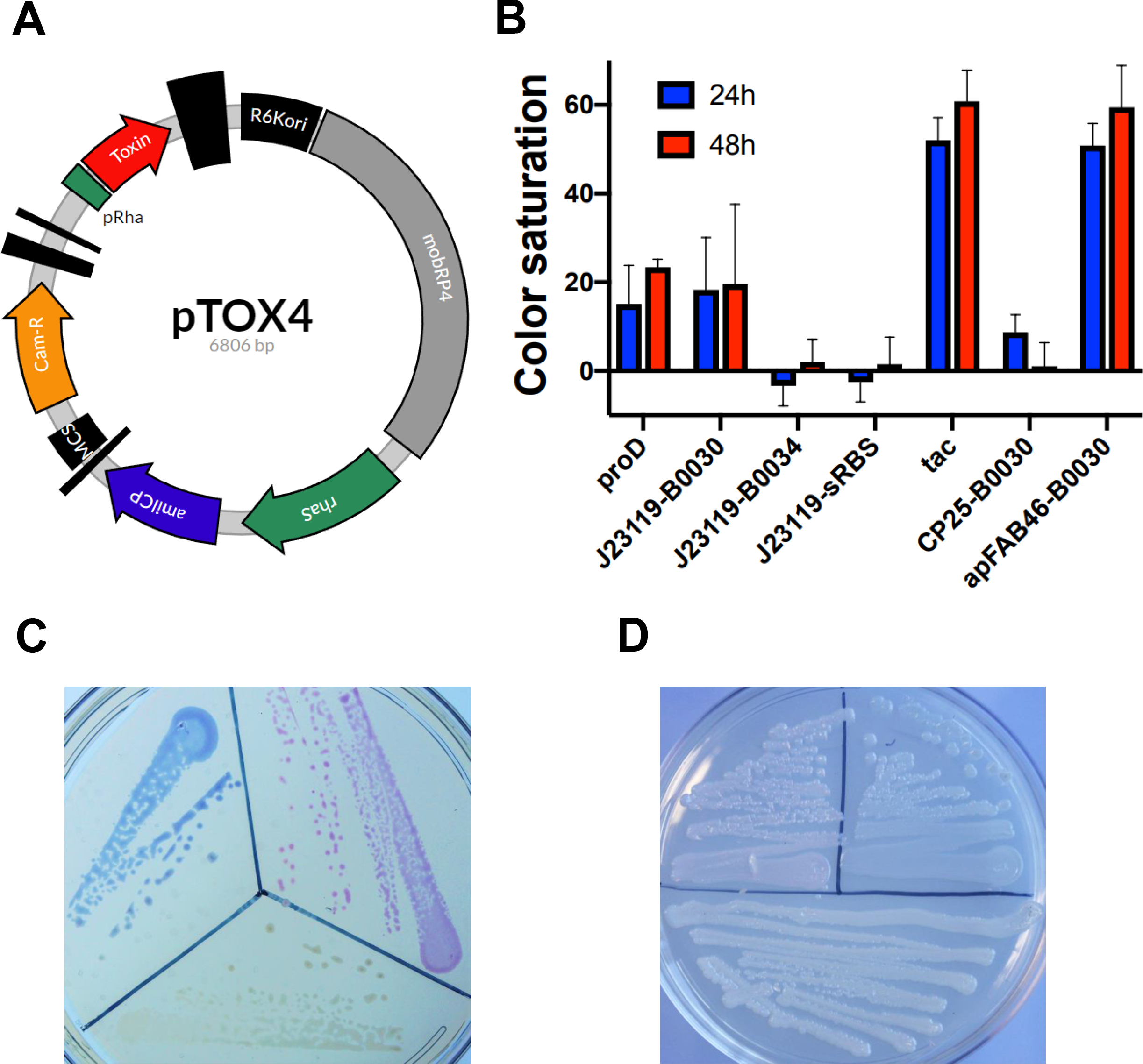
A chromoprotein module facilitates monitoring of conjugation. A) Plasmid map of pTOX4. R6Kori, the R6K origin of replication; mobRP4, mobilization region from RP4 conjugative plasmid; *rhaS*, the rhamnose transcriptional activator; *amilCP*, the blue amilCP chromoprotein; MCS, multiple cloning site; Cam-R, chloramphenicol resistance cassette; pRha, rhamnose promoter. Vertical black bars of varying width represent terminators. B) tac and apFAB46-B0030 allow optimal amilCP expression. Relative color saturation at 24h and at 48h of pTOX4-containing colonies with various promoters and ribosome-binding sites (described in more detail in the Methods). C) Depiction of donor *E. coli* containing (from bottom, clockwise) pTOX without chromoprotein, with tac-*amilCP*, and with apFAB46-B0030-*tsPurple* after 24h at 37°C. D) *E. cloacae* pTOX merodiploids (from bottom, clockwise) without chromoprotein, with tac-*amilCP*, and with apFAB46-B0030-*tsPurple* after 24h at 37°C and an additional 24 hours at 25°C.

The series of *amilCP* modules were first tested in *E. coli* donors, where we found that the tac promoter (21) or the apFAB46 promoter (22) offered the deepest blue coloration (Fig 4B, Fig S3A). This level of *amilCP* expression did not incur a detectable fitness cost (Fig S3B); however, several strategies to increase colony coloration further (e.g increasing *amilCP* copy number) led to toxicity and were not pursued. pTOX vectors containing *amilCP* driven by the tac promoter and a pTOX vector expressing the magenta *tsPurple* chromoprotein driven by apFAB46 were created (Fig 4C) (23). Both AmilCP and TsPurple were visible in re-streaked merodiploid colonies after 24-48 hours of incubation (Fig 4D), though coloration was not as saturated as when expressed from the pTOX plasmids (which have medium-copy origins). Therefore, the pTOX chromoprotein modules may prove useful for monitoring the success of single- and double-crossover, particularly in organisms with inefficient conjugation.

### Application of the pTOX vectors to study inducible antibiotic resistance

To further interrogate the utility of the pTOX suite, we used these vectors to characterize the role of the *S. marcescens* peptidoglycan (PG) amidohydrolases in inducible beta-lactam resistance mediated by the AmpC beta-lactamase. The PG component of the bacterial cell wall consists of a repeated disaccharide polymer linked through peptide cross-links. The peptidoglycan amidohydrolases facilitate remodeling of the cell wall by catalyzing hydrolysis of the amide bond linking the polysaccharide to the peptide component, generating muropeptide breakdown products that can subsequently be recycled in the cytoplasm (24). When the classical cytoplasmic PG amidohydrolase, *ampD,* becomes saturated with substrate in the setting of catastrophic remodeling precipitated by beta-lactam antibiotics such as penicillins and cephalosporins, the accumulation of muropeptides leads to *ampC* derepression. The associated beta-lactam resistance enables subsequent restoration of PG homeostasis (25).

In *E. cloacae* and *Citrobacter freundii,* expression of *ampC* at basal levels is sufficient for clinical resistance to penicillins and early-generation cephalosporins. After exposure to beta-lactams and the resulting accumulation of muropeptide breakdown products, transcriptional upregulation can lead to transient intermediate resistance to late-generation cephalosporins such as ceftriaxone. Under conditions where there is selection for high-level cephalosporin resistance (i.e. in individual patients who are subjected to prolonged cephalosporin treatment), mutation of the *ampD* amidohydrolase can occur. This leads to a constitutive increase in cytoplasmic muropeptide that is sufficient for high level derepression of *ampC* and resistance to ceftriaxone (26, 27). However, it is unclear whether the insights gained from studies of *E. cloacae* and *C. freundii* can be generalized to all AmpC-producing organisms, because the pathway to full derepression may be more complicated in organisms with multiple orthologous amidohydrolases. For example, in *Pseudomonas aeruginosa*, full derepression of *ampC* requires inactivation of additional *ampD* orthologues (28), while in *Yersinia enterocolitica*, deletion of all three *ampD* orthologues does not result in obvious clinical resistance (29).

Systematic investigation of the contribution of *S. marcescens* PG amidohydrolases to *ampC* derepression and resulting beta-lactam resistance has not been performed. We found that *S. marcescens* encodes 3 PG amidohydrolases, which, by sequence homology (30) we denote *ampD* (WP_033641266.1), *amiD* (WP_016928349.1), and *amiD2* (WP_048796451.1) (Fig 5A, Fig S4). Creation of pTOX derivatives targeting each of the *S. marcescens* PG amidohydrolases allowed the rapid generation of all combinations of single, double, and triple mutants (Fig. S5). We found that, of the single mutants, only Δ*amiD2* had a significant increase in basal AmpC activity (Fig. 5B); however, this corresponded to only a 2-fold increase in cephalosporin MICs (Table 2). In contrast, the Δ*ampD*Δ*amiD2* double mutant had a more than 50-fold increase in AmpC activity, which resulted in an 8-fold increase in the ceftriaxone and a 4-fold increase in cefepime MICs. The triple mutant exhibited no further increase in AmpC activity or in MICs. The Clinical and Laboratory Standards Institute (CLSI) has recently updated their guidelines on MIC breakpoints above which there is a potential for clinical resistance. Under the new breakpoints, the Δ*ampD*Δ*amiD2* double mutant and triple mutant, with MICs of 2, would be considered to have “intermediate” resistance to ceftriaxone, but to still be fully sensitive to ceftazidime and cefepime. In comparison, inactivation of the single *E. cloacae ampD* was reported to result in a ceftriaxone MIC of 32 (from a baseline of 0.5) (26).

**TABLE 2.**
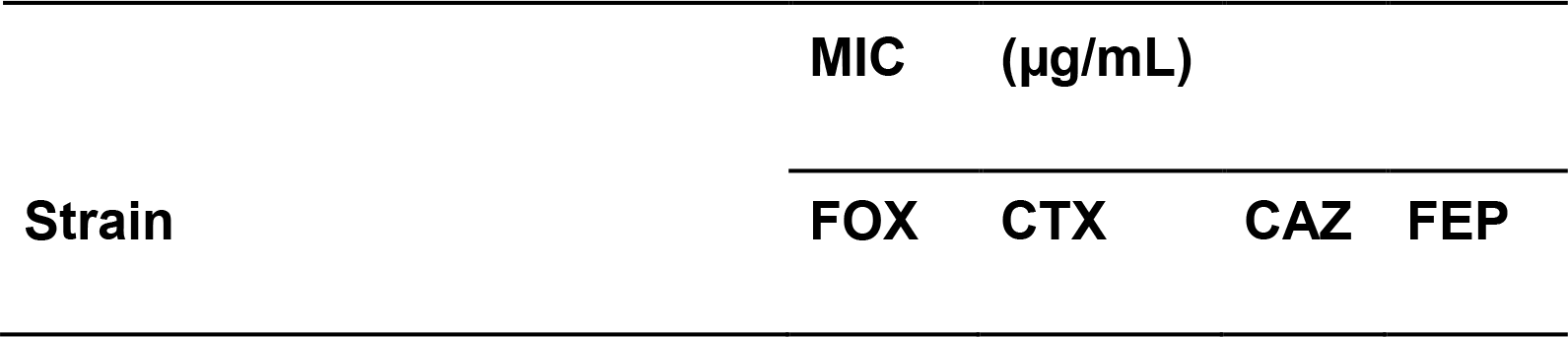
Minimal inhibitory concentrations for amidohydrolase mutants^b^

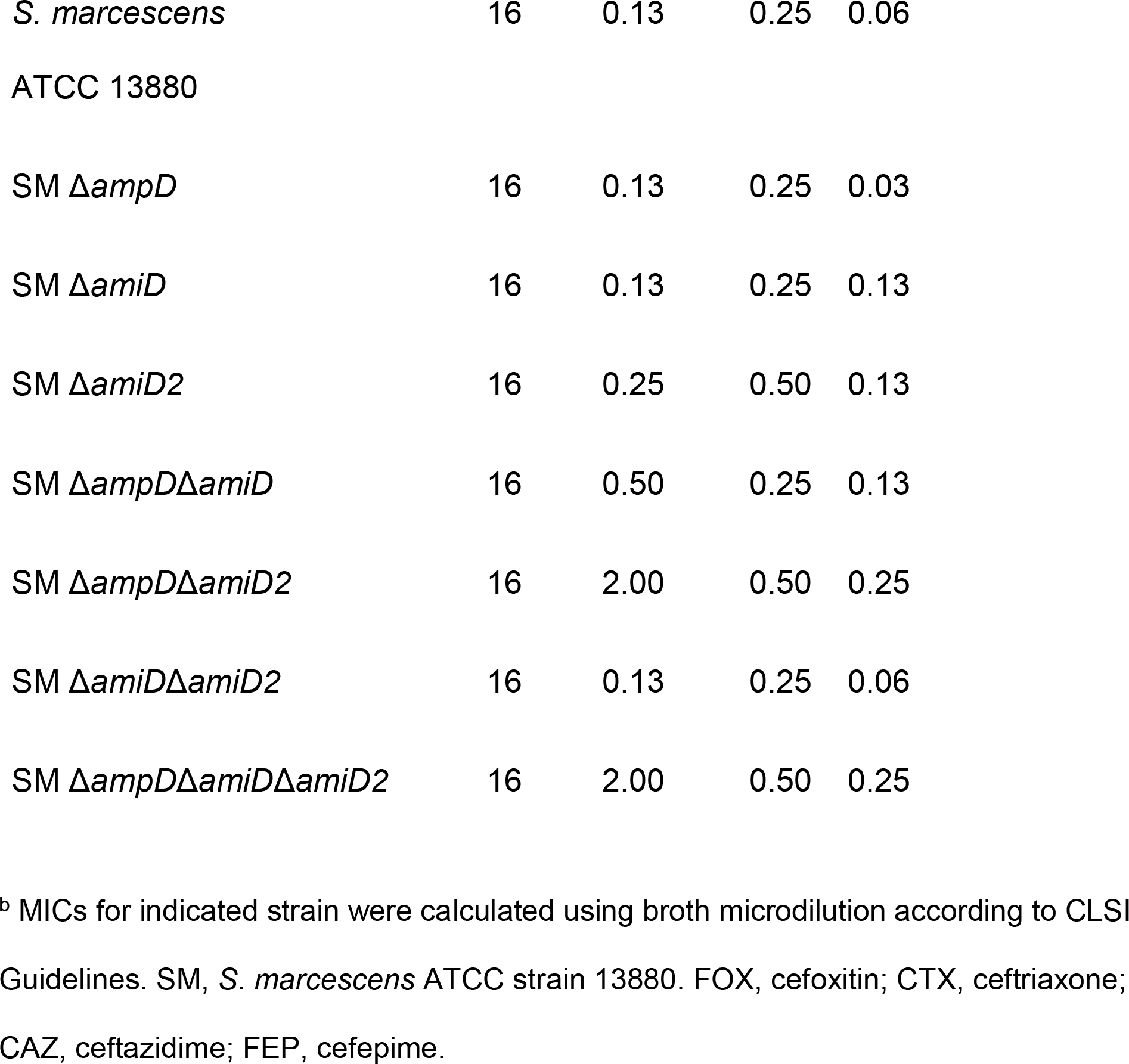

**Fig 5.**
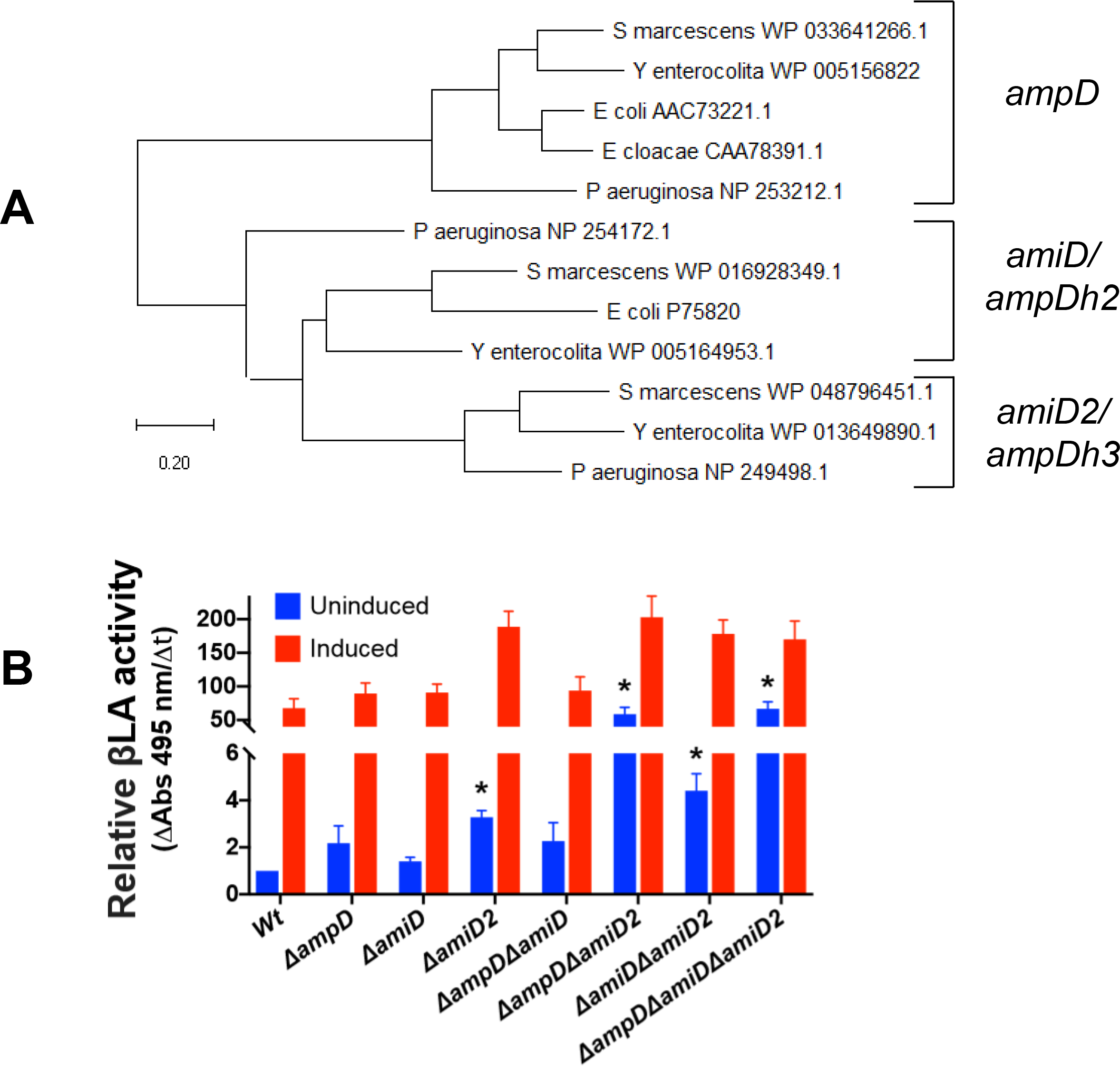
*S. marcescens* peptidoglycan amidohydrolase deletions lead to differential derepression of *ampC.* A) Phylogenetic analysis performed using the Maximum Likelihood method and JTT matrix-based model in MEGA X (30)An unrooted tree is shown with the lowest log likelihood (−4913). B) Clarified sonicates from indicated strains were incubated with equal amounts of nitrocefin, a chromogenic cephalosporin beta-lactam, and absorbance measured in kinetic mode for 10 minutes. The slope of the line from the first 5 data points were used to calculate beta-lactamase activity, which was then normalized to Wt. Measurements are shown either without pre-induction, and those with induction with cefoxitin 4 μg/mL for 2 hours prior to harvesting. Data represent the mean ± SEM of 4 independent experiments. Comparisons were made between all uninduced mutants and Wt, and between each induced sample and its uninduced control. * = p < 0.05 after performing Bonferroni correction. All induced samples are also significantly different from their uninduced samples, except for Δ*ampD*Δ*amiD*, Δ*ampD*Δ*amiD2*, and the triple mutant. These asterisks are not shown for clarity.

## Discussion

Our findings suggest that the pTOX suite of allele exchange vectors described here should facilitate the genetic engineering of diverse Proteobacteria. Each of the pTOX vectors includes a rhamnose inducible toxin that may circumvent escape from counter selection, which can limit *sacB*-based allele exchange vectors. These toxins have been used to facilitate recombineering (11), and inducible toxins for allelic exchange promise to be a broadly generalizable approach, as systems have recently also been described for Vibrio and Aeromonas species (31) as well as for the archaeon *Pyrococcus yayanosii* (32).

The pTOX vectors contain an expanded multiple cloning site, multiple antibiotic resistance cassettes, and chromoprotein modules that facilitate monitoring of crossover events. The utility of the pTOX vectors and all 3 of the different toxins they encode was demonstrated through creation of multiple deletions in 4 different pathogens, including *S. flexneri*, an organism in which allele exchange has been difficult. All of these vectors have been deposited at Addgene to facilitate their distribution. Besides their utility for engineering Gram-negative organisms in research labs, these vectors may also be useful in the context of undergraduate education.

*S. marcescens,* along with *E. cloacae*, *C. freundii, Klebsiella aerogenes* and *Morganella morganii*, are members of a group of pathogenic *Enterobacteriaceae* with the potential for high level, inducible expression of AmpC, which in some cases has been linked to resistance to almost all penicillins and cephalosporins (33). Original reports of cephalosporin failure in *E. cloacae* (34) engendered the practice of using ultra-broad spectrum antibiotics (such as cefepime or carbapenems, which are resistant to AmpC hydrolysis) in the treatment of serious infections by pathogens with the potential for AmpC overexpression. However, this approach has untoward consequences, including increasing infections with carbapenem-resistant Enterobacteriaceae (35).

It is not clear if routine use of ultra-broad spectrum antibiotics is warranted for all organisms with inducible AmpC expression. A recent review (36) emphasized that besides *E. cloacae,* the data on cephalosporin failure for pathogens with inducible AmpC is sparse. What data do exist emphasize that true on-treatment emergence of beta-lactam resistance is probably rare, at least in *S. marcescens* and in *Morganella morganii (37)*. *In vitro* experiments also hint at important heterogeneity among these pathogens; in this setting, the development of spontaneous cephalosporin resistance has been reported to be nearly 100-fold lower in *S. marcescens* compared to *E. cloacae* and *C. freundii*, and 10-fold lower still in *M. morganii* (38).

Our observations suggest that ultra-broad-spectrum antibiotics may be not be necessary for treatment of *S. marcescens* infections. We used the pTOX vectors to investigate the role of *S*. *marcescens’* 3 peptidoglycan amidohydrolases on inducible beta-lactam antibiotic resistance. We found that deletion of a single amidohydrolase locus had a minimal effect on cephalosporin MICs, and that even the absence of all 3 amidohydrolase loci did not consistently render *S*. *marcescens* resistant to this class of antibiotics, although the triple mutant and the *ΔampDΔamiD2* double mutant did exhibit intermediate resistance to ceftriaxone. Thus, the effects of amidohydrolase deletion in *S. marcescens* differ from those in *C. freundii* and *E. cloacae,* in which resistance arises following the loss of a single amidohydrolase. Importantly, though current CLSI breakpoints would classify the *ΔampDΔamiD2* double mutant as having “Intermediate” resistance to ceftriaxone, there is no evidence of increased clinical failure in this range (39). This is important since ceftriaxone is less expensive, has more convenient dosing intervals, and is a narrower spectrum agent compared to cefepime or carbapenems. Further work with additional *S. marcescens* isolates to clarify the generalizability of our findings is warranted.

## Materials and Methods

### pTOX construction

The DNA components for the pTOX series were obtained from pDS132 (15), the pSLC recombineering series (11) which was a gift from Swaine Chen (Addgene plasmid # 73194), pON.mCherry (21) which was a gift from Howard Shuman (Addgene plasmid # 84821), strain TP997 (40) which was a gift from Anthony Poteete (Addgene plasmid # 13055), and direct gene synthesis (from Integrated DNA Technologies) and were assembled using Gibson or HiFi Assembly (New England BioLabs) unless otherwise stated. All restriction enzymes were obtained from New England Biolabs and all PCR was performed with primers from Integrated DNA Technologies and Q5 polymerase (New England Biolabs). All cloning steps were performed in π-carrying hosts (either DH5αpir (41) for propagation or MFD-π (42) for conjugation) under catabolite repression in LB containing the appropriate antibiotic and 2% glucose (w/v).

pSLC toxin vectors were first linearized with primers prJL1 and prJL2 and joined with the fragment obtained from pDS132 with primers prJL3 and prJL4 (see Supplemental Table 1 for all primers used in this study). The mobRP4 from pDS132 was subsequently amplified with primers prJL5 and prJL6 and assembled with the prior vectors cut with NheI. The chloramphenicol resistance cassette from pON.mCherry was then amplified with primers prJL7 and prJL8 and inserted into the prior vectors digested with ClaI and BglII. The π-dependent origin from pDS132 was next isolated by SmaI-digestion and inserted into the prior vectors linearized with prJL9 and prJL10. A codon-optimized *rhaS* (with the original primary protein sequence obtained from WP_000217135.1) and promoter (see Supplemental Text 1 for the sequence of all directly synthesized DNA fragments used in this study) was obtained by direct synthesis and assembled into the prior vectors linearized with prJL11 and prJL12. The expanded polylinker (19) with the forward transcriptional terminator BBa_B1002 (IGEM) was obtained by direct synthesis (Sequence 2) and inserted into the prior vectors linearized with primers prJL13 and prJL14. The artificial ribosome binding site was generated using the online calculator derived after (17), synthesized as above (Sequence 3) and assembled into the prior vectors linearized with prJL15 and prJL16 to generate pTOX1 (containing *yhaV*), pTOX2 (containing *mqsR*) and pTOX3 (containing *tse2)*. See Supplemental Table 2 for all plasmids used in this work. For insertion of *amilCP* or *tsPurple*, the above vectors were cut with SbfI and Sequence 4 and Sequence 5 inserted. For replacement of the chloramphenicol resistance cassette with one encoding gentamicin resistance, the appropriate vector was linearized with prJL17 and prJL18 and assembled with the cassette amplified from strain TP997 (using prJL19 and prJL20). See Supplemental Table 3 for all strains used in this work. Q5 GC enhancer (New England Biolabs) was used for amplification of mobRP4 and *tse2*.

### Insertion of homology targeting regions

pTOX vectors were cut with SmaI and the relevant homologous regions were assembled after being amplified with prJL21, prJL22, prJL23, and prJL24 (for *S. marcescens hexS);* prAW1, prAW2, prAW3, and prAW4 (for EHEC *lacZ)*; prCJK1, prCJK2, prCJK3, and prCJK4 (for *S. flexneri ipgH);* prJL25, prJL26, prJL27, and prJL28 (for *E. cloacae ampC)*; prJL29, prJL30, prJL31, and prJL32 (for *S. marcescens ampD*); prJL33, prJL34, prJL35, and prJL36 (for *S. marcescens amiD*); and prJL37, prJL38, prJL39, and prJL40 (for *S. marcescens amiD2*). Note that some of the overlap regions in the above primers correspond to a version of pTOX with the original pDS132 polylinker.

### Allelic exchange with pTOX

On day 1, the appropriate upstream and downstream sequences from the targeted pathogen are amplified from gDNA in separate PCR reactions. After column purification of the resulting PCR product (Denville), the products are assembled with pTOX previously gel-purified after restriction digestion of the polylinker and electroporated into an *E. coli* strain that could serve as donor in conjugation. Throughout this work, we routinely used the diaminopimelic acid (DAP) auxotroph MFD-π (42) as the pTOX donor strain. Unless specified, all subsequent steps are performed in the presence of 2% glucose to avoid premature toxin induction. On day 2, colony PCR was performed on single MFD-π transformant colonies to confirm the appropriate insert size. On day 3, conjugation was performed between the MFD-π bearing pTOX and the pathogen of interest. Optimizing the conjugation is crucial. For example, we found that conjugation was efficient at 4-8 hours at 37°C with a 3:1 (v/v) ratio of MFD-π:pathogen for EHEC, *E. cloacae*, and *S. flexneri,* but *S. marcescens* had markedly better efficiency when conjugated overnight at 30°C using 50-fold excess volume of an early logarithmic phase growth culture of MFD-π. Exconjugants were isolated on appropriate antibiotics. On day 4, a single exconjugant colony is resuspended in 2 mL of LB containing glucose (but no selective antibiotic). This culture is incubated at 37°C with agitation until OD600 0.2, then washed twice with M9 salts (Sigma) with 2% rhamnose (w/v) before plating on the M9-rhamnose agar described below. A short preliminary outgrowth in broth without selection minimizes the possibility of the culture becoming dominated with a single double-crossover rhamnose-resistant clone. On day 5, the desired mutants can be identified with colony PCR on the resulting double-crossover colonies. The selection is stringent and in this manner, individual colonies can frequently be isolated from a plate inoculated with the undiluted washed culture from above, but 10^−1^ and 10^−2^ dilutions should also be plated.

For the experiments described in Table 1, primers prJL51 and prJL52 were used; their amplicon consisted of a small intergenic region that was largely replaced when the expanded polylinker was inserted.

### *amilCP* coloration optimization

pTOX derivatives with different promoters and ribosome binding sites to drive *amilCP* were created by assembling SbfI-cut pTOX1 with *amilCP* (Sequence 4) amplified with prJL59 and either prJL41 (for J23119-B0030), prJL42 (for J23119-B0034), prJL43 (for CP25-B0030), or prJL44 (for apFAB46-B0030). The J23119 promoter and B0030 and B0034 ribosome binding sites sequences were obtained from IGEM. The insulated proD promoter (43) was amplified from pSB3C5-proD-B0032-E0051 (which was a gift from Joseph Davis and Robert Sauer; Addgene plasmid #107241) with prJL47 and prJL48, fused by SOE PCR with the *amilCP* coding sequence obtained from PGR-Blue (44) (which was a gift from Nathan Reyna; Addgene plasmid #68374) using prJL49 and prJL50, and after XbaI-digestion of this product, it was ligated with XbaI-cut pTOX1. The J23119-synthetic ribosome binding site (17) was amplified from Sequence 6 with prJL45 and prJL46 and assembled in SbfI-cut vector and *amilCP* amplified with prJL49 and prJL50 as for proD above.

*E. coli* DH5αpir were transformed with the appropriate *amilCP*-containing plasmid. Single colonies were grown in overnight cultures, diluted 1:100, and then back-diluted once in logarithmic phase growth so to enable spot-streaking onto solid agar at the identical optical density. Digital images were taken at 24h and 48h and saturation obtained by splitting the resulting image into an “HSB Stack” in ImageJ. The peak saturation was subsequently obtained using the “Measure” function, then normalized by subtracting the peak saturation of the resulting spots from spots of *E. coli* DH5αpir carrying pTOX1 without *amilCP.* The resulting values represent the mean ± SEM of this procedure done on 3 different days.

### Beta-lactamase assay

Overnight cultures of indicated strains were back-diluted 1:100 (v/v) into fresh media and grown for an additional 2 hours. Bacteria were then pelleted, washed twice in phosphate-buffered saline, and then flash-frozen in liquid nitrogen. On the day of the assay, pellets were thawed at 37°C and then subjected to a single round of sonication on ice (Sonic Dismembranator 60, Fisher Scientific, setting 8, 5 seconds). Lysates were clarified by centrifugation at 20,000 rcf for 60 minutes at 4°C. Total protein was quantitated by fluorometry using the Qbit Protein Assay Kit (Thermo Fisher). Beta-lactamase activity was determined by the addition of 80 ng nitrocefin to either 250 ng or 1000 ng of total protein; to facilitate accurate quantitation, 250 ng was used for all cefoxitin-induced *S. marcescens* samples and also for the Δ*ampD*Δ*amiD2* double mutant and the triple mutant. Immediately after addition of nitrocefin with a multi-channel pipettor, absorbance was read kinetically at 495 nm every 5 minutes in a Synergy HT plate reader (BioTek). For Figure 5, the slope of the absorbance was normalized to wild-type *S. marcescens* and the amount of total protein added.

### Minimum inhibitory concentration (MIC)_determination

Minimum inhibitory concentrations were determined for the indicated *S. marcescens* isolates and performed by broth microdilution according to CLSI guidelines and after Weigand *et al (45)*. Briefly, overnight cultures were back-diluted in cation-adjusted Mueller-Hinton broth, allowed to grow for 2 hours, and adjusted to a final inoculum of 5 x 10^5^ colony-forming units per mL before applying to wells with the appropriate antibiotic concentration. Results were read after 20 hours of incubation at 37°C. The results in Table 2 represent the mode of 3 independent experiments.

### Materials and strains

Unless otherwise specified, all materials were purchased from Sigma. When appropriate, media was supplemented with streptomycin 200 µg/mL, gentamicin 5 µg/mL, and chloramphenicol 20 µg/mL for all *E. coli*, *E. cloacae,* and *S. flexneri. S. marcescens* exconjugants were isolated at 100 µg/mL chloramphenicol. Diaminopimelic acid (DAP) was used at a final concentration of 0.3 mM, 5-bromo-4-chloro-3-indolyl-β-D-galactopyranoside (X-gal) at 60 µg/mL, glucose at 2% (w/v) in all propagation steps with pTOX vectors. When washing the out-grown single-crossovers, rhamnose was used at 2% (w/v) in M9 salts. The resulting washed bacteria were plated on M9 agar supplemented with 0.2% casamino acids (w/v), 0.5 mM MgSO_4_, 0.1 mM CaCl_2_, 25 uM iron chloride in 50 uM citric acid, the appropriate antibiotic, and rhamnose. Rhamnose at a final concentration of 0.2%-2% facilitated good toxin induction in the organisms we tested; there was no obvious correlation with the concentration of rhamnose used, but it may be prudent to optimize this in new organisms. The *S. marcescens* ATCC 13880 isolate used throughout this work is a spontaneous mutant selected on streptomycin. *E. cloacae* was obtained from ATCC (isolate 13047). EHEC was isolate EDL933. *S. flexneri* was strain 2457T.

### Miscellaneous analysis

All figures and statistical analyses were prepared in Prism 8 (Graphpad). The growth curves in Supplemental Figure 3 were generated using Bioscreen C (Growth Curves USA). The plasmid maps were generated with AngularPlasmid and ApE (for the polylinker inset in Figure 1).

## Acknowledgments

JEL has been supported by T32 AI-007061 and by the Harvard Catalyst Medical Research Investigator Training fellowship; ARW by T32AI132120; MKW by R01 AI-042347 and HHMI. We thank the other members of our group for many productive conversations informing the design of pTOX and for comments on the manuscript.

---

**Fig S1.**
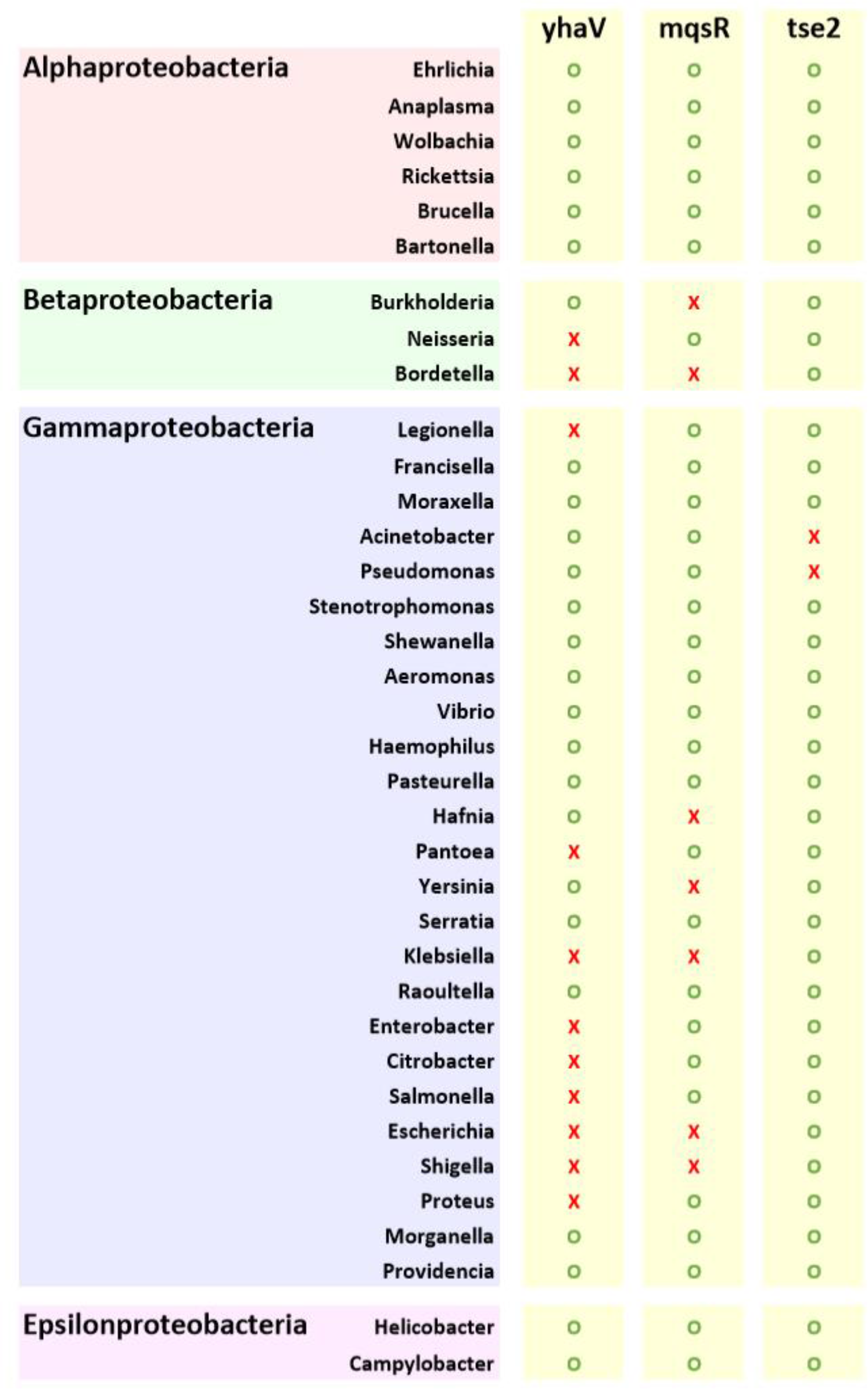
Toxin(s) predicted to be useful (green open circle) in diverse pathogens based on absence of toxin homolog by BLASTP in high confidence genomes deposited in NCBI. Red X’s denote presence of toxin gene.

**Fig S2.**
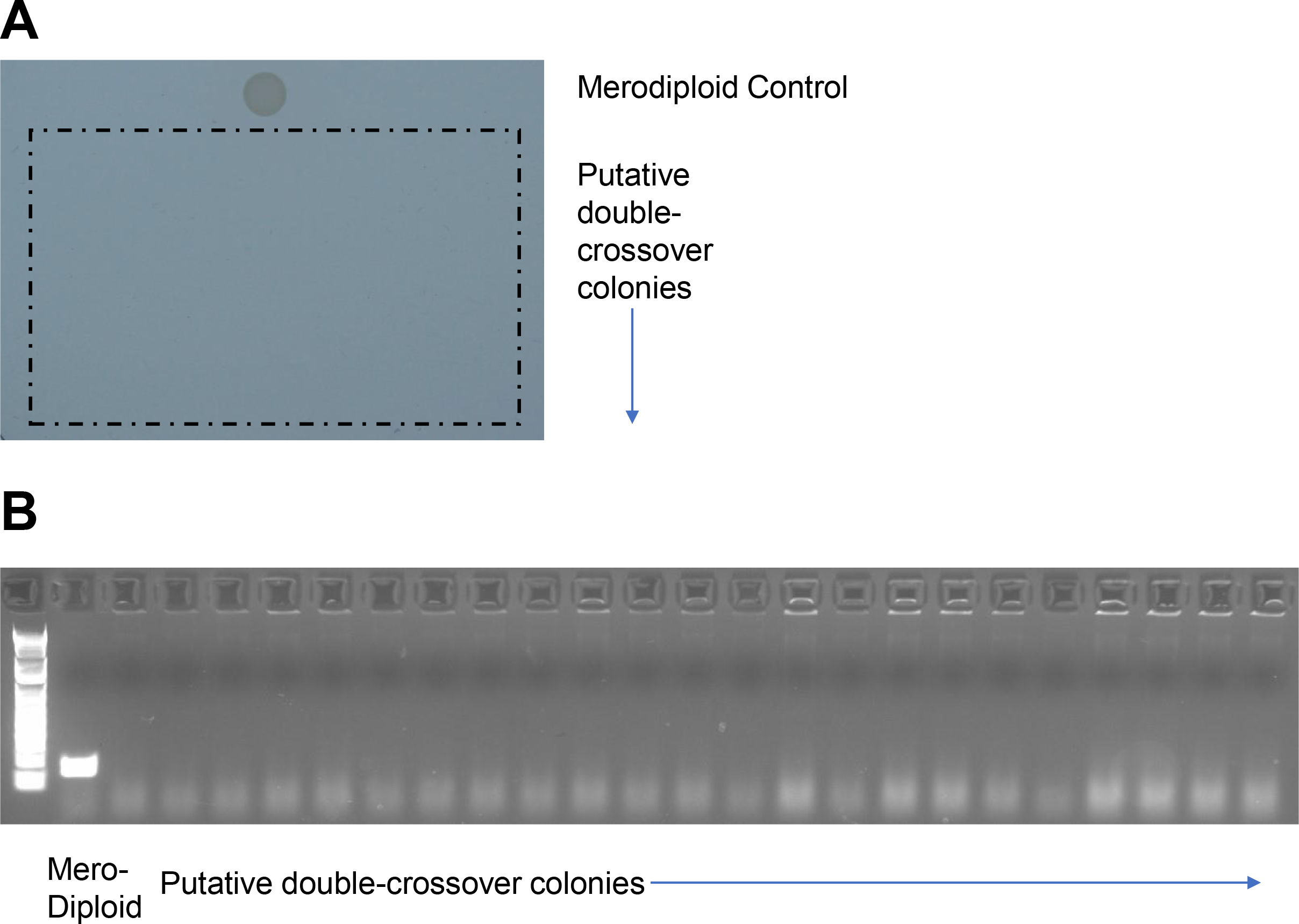
Putative double-crossover *S. marcescens* colonies (for each toxin, 23 colonies from 3 different crossovers) were randomly selected for characterization. Summary results are in Table 1. A) A colony was resuspended in non-selective LB and then spotted onto LB+chloramphenicol. The top colony is the merodiploid positive control. There is no growth where the putative double-crossover colonies were spotted (hatched box). B) Colony PCR was performed for a small intergenic amplicon (between *rhaS* and the chloramphenicol resistance promoter) to test for presence of retained pTOX. The first lane is the merodiploid positive control.

**Fig S3.**
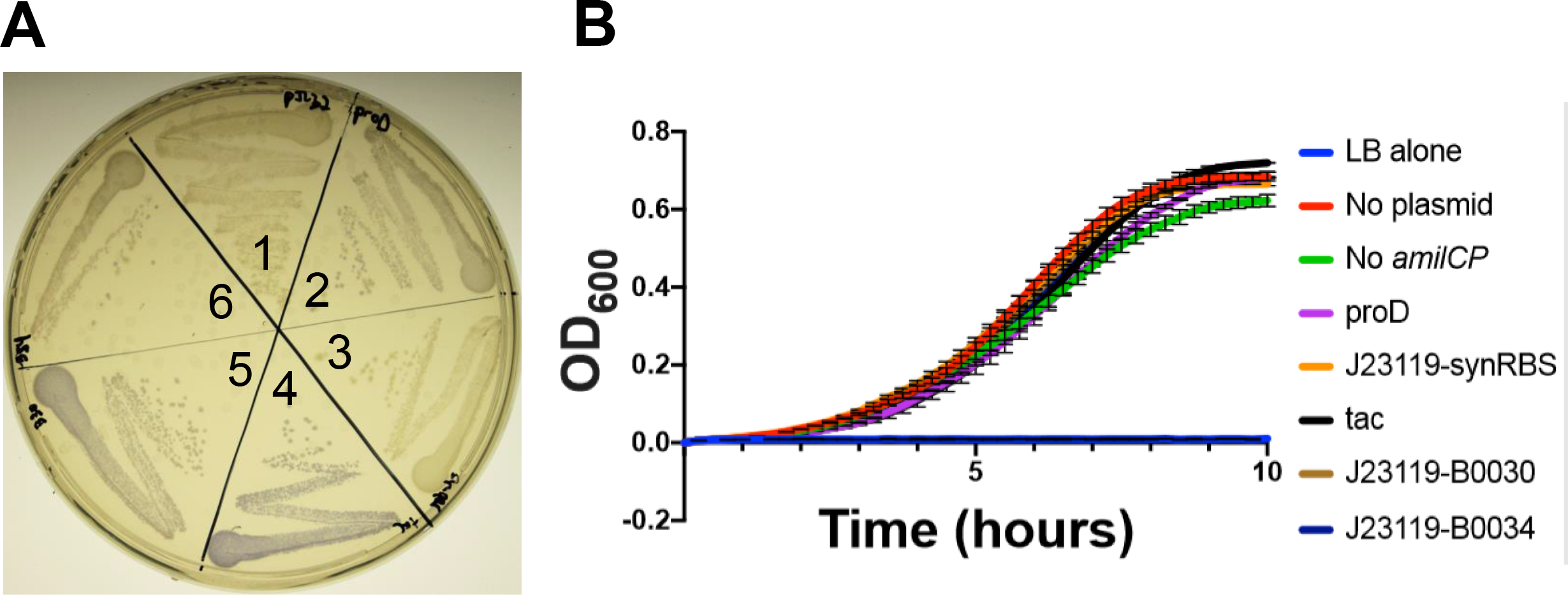
A) *E.* coli donors containing pTOX vectors with *amilCP* and various promoters and ribosome binding sites (RBS) at 24h at 37°C. From top, clockwise: 1) pTOX with no *amilCP;* 2) pTOX-*amilCP* with proD promoter; 3) pTOX-*amilCP* with J23119 promoter and synthetic RBS; 4) pTOX-*amilCP* with tac promoter; 5) and 6) pTOX-*amilCP* with J23119 promoter with the B0030 or B0034 RBS, respectively. B) Growth curves for *E. coli* donor strains (or LB alone) containing pTOX with *amilCP* driven by indicated promoter/RBS.

**Fig S4.**
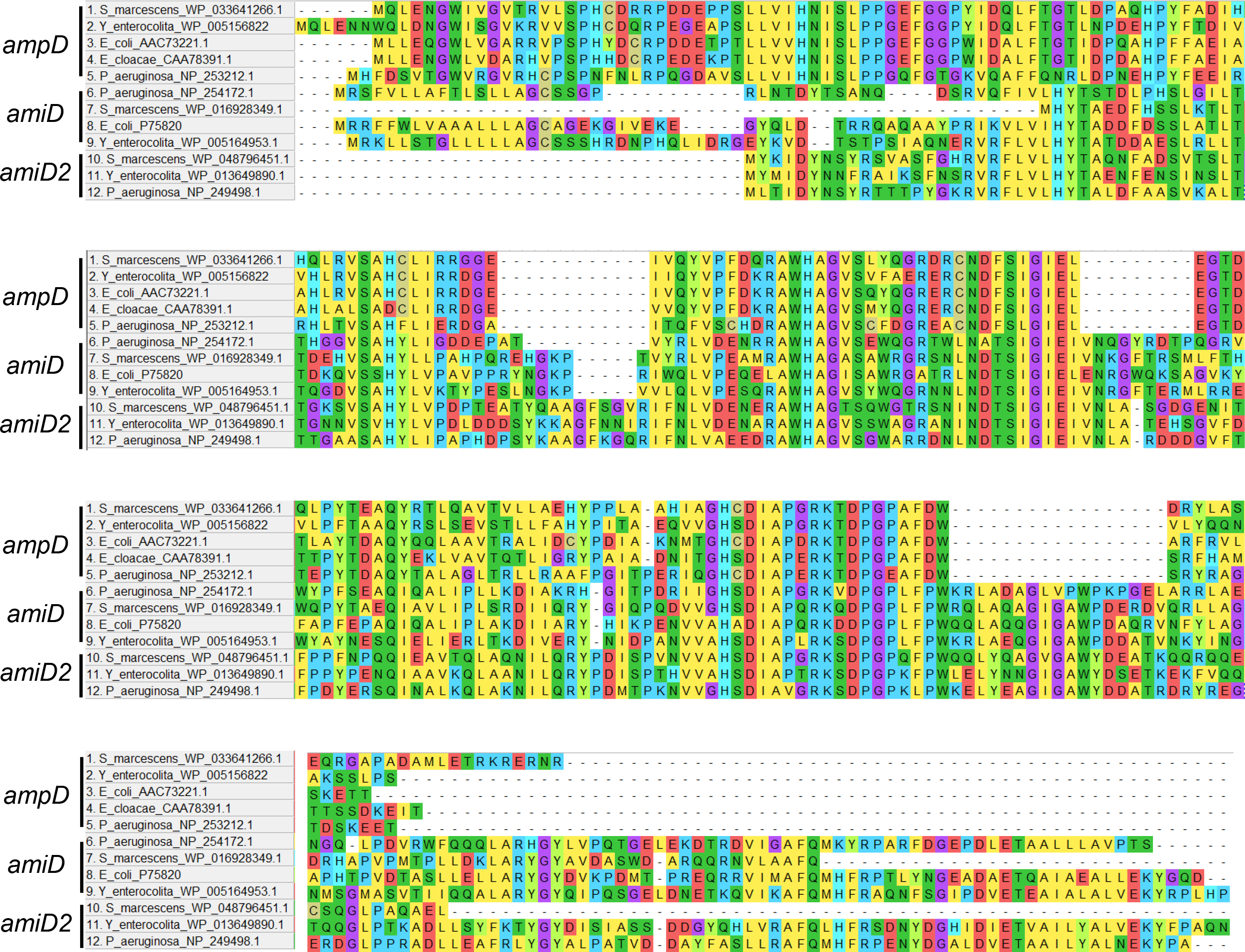
Indicated sequences were aligned in MEGA X using the MUSCLE algorithm (with the UPGMA method). The results are depicted as sequential (reading from left to right from the first through fourth panel) primary sequence.

**Fig S5.**
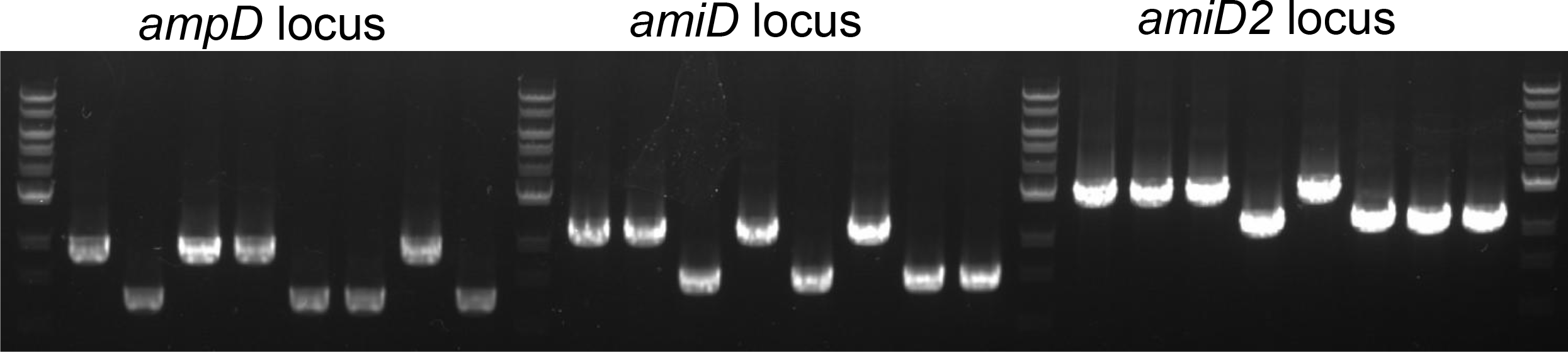
PCR-based genotyping of *S. marcescens* amidohydrolase mutations. Lanes from left to right: marker, Wt, Δ*ampD,* Δ*amiD,* Δ*amiD2,* Δ*ampD*Δ*amiD,* Δ*ampD*Δ*amiD2,* Δ*amiD*Δ*amiD2,* Δ*ampD*Δ*amiD*Δ*amiD2.* The *ampD* locus was amplified with prJL53 and prJL54; *amiD* with prJL55 and prJL56; and *amiD2* with prJL57 and prJL58

## Supplemental Text 1

### Sequence 1 - Codon-optimized *rhaS* with promoter

cgcggaacccctatttgtttatttttctaaatacattcaaatatgtatccgctcatgagacaataaccctgataaatgcttcaataatattgaaaaaggaagagtATGACGGTGCTGCACTCGGTTGACTTCTTCCCTAGCGGCAATGCCAGCGTTGCCATTGAGCCGCGCCTGCCTCAAGCCGACTTCCCGGAGCACCACCACGACTTCCACGAGATCGTTATCGTGGAGCACGGTACCGGCATCCACGTTTTCAACGGCCAACCGTACACGATTACGGGCGGTACCGTGTGCTTCGTTCGTGATCACGACCGCCACTTATACGAGCACACGGACAACTTATGCTTAACCAACGTTTTATACCGTAGCCCTGACCGCTTCCAATTCCTGGCGGGCTTAAACCAACTGTTACCGCAGGAATTAGACGGCCAATACCCTAGCCATTGGCGTGTGAATCATTCGGTGCTGCAACAAGTTCGCCAATTAGTGGCGCAAATGGAGCAACAAGAGGGCGAGAACGACCTGCCGAGCACGGCGAGCCGTGAAATTCTGTTCATGCAGCTGTTACTGCTGTTACGCAAGTCGAGCCTGCAAGAAAATTTAGAGAATTCGGCGAGCCGCCTGAATCTGCTGTTAGCGTGGTTAGAAGATCACTTCGCGGACGAAGTTAACTGGGATGCGGTTGCCGACCAGTTCAGCCTGAGCTTACGCACCCTGCACCGCCAACTGAAACAACAGACCGGCTTAACCCCGCAACGCTATTTAAACCGTTTACGCTTAATGAAGGCGCGCCACTTACTGCGCCATTCGGAAGCGTCGGTGACCGATATTGCGTACCACTGTGGCTTTTCGGATAGCAATCATTTCAGCACCCTGTTCCGTCGCGAATTCAATTGGAGCCCTCGCGACATCCGCCAAGGCCGCGACGGTTTCTTACAGTGA

### Sequence 2 - Forward terminator and expanded polylinker

AGCTTCGAGCTAATCGcgcaaaaaaccccgcttcggcggggttttttcgcTGATCACGTACGATATCTTCGAACCGGTGCACATGGTGTACAGGGCCCTAGGATAGGACGTCTTAAGGTTTAAACCAGGTTAATTAATTTAAATGCATCCCGGGACGTCTCGAGCTCGATCGGACCGCGGCCGCTAGCACGTATACCAAGTGTCCTGT

### Sequence 3 - Synthetic strong ribosome binding site

ttatttttctaaatacattcaaatatgtatccgctcatgagacaataaccctgataaatgcttcaataatattgaaaaaggaGCTTATCACCGATAAGGAGGTTTTTTAATGACGGTGCTGCACTCGGTTGACTTCTTCCCTAGCGGCAATGCCAGCGTTGCCATTGAGCCGCGCCTGCCTCAAGCCGACTTCCCGGAGCAC

### Sequence 4 - *amilCP* with tac promoter

ttgacaattaatcatcggctcgtataatgtgtggaaaggcggttcaccgccgttttcacacaggaaacagaattctttaagaaggagatatacatATGTCAGTGATAGCAAAGCAGATGACATACAAAGTATATATGAGCGGTACTGTAAATGGTCACTACTTCGAAGTAGAAGGTGATGGCAAAGGGAAGCCGTATGAGGGTGAACAAACAGTGAAGCTTACAGTTACGAAGGGTGGCCCTTTGCCGTTCGCGTGGGACATTTTGTCGCCACAGTGCCAGTACGGGAGCATACCATTCACCAAGTATCCAGAGGACATACCAGACTACGTGAAACAGTCCTTTCCCGAAGGCTATACCTGGGAGCGCATAATGAACTTTGAAGACGGTGCGGTTTGTACGGTATCGAACGATTCATCAATCCAAGGAAATTGCTTTATTTATCATGTTAAATTTTCGGGGCTTAACTTTCCGCCAAATGGCCCCGTGATGCAGAAGAAAACTCAGGGCTGGGAACCAAATACCGAGCGCCTTTTCGCTCGGGACGGAATGCTGTTGGGAAACAATTTTATGGCGTTGAAGCTGGAAGGGGGTGGCCACTATCTCTGTGAATTCAAGACAACATACAAAGCCAAGAAGCCTGTCAAGATGCCCGGTTATCACTATGTGGACCGGAAGCTCGATGTCACTAATCACAACAAGGATTATACTTCAGTTGAACAGTGCGAAATTTCGATCGCTCGCAAACCTGTCGTAGCATAA

### Sequence 5 - *tsPurple* with apFAB46 promoter

AGGCCTtctagagtcgacctgcaaaaaagagtattgacttcgcatctttttgtacctataatagattcattactagagaaagaggagaaatactagATGGCGTCCCTTGTAAAGAAGGATATGTGTGTTAAGATGACAATGGAGGGAACCGTCAACGGGTATCACTTTAAGTGTGTTGGCGAGGGAGAAGGCAAACCGTTCGAGGGTACACAGAATATGCGCATTCGTGTCACCGAAGGGGGCCCTTTGCCCTTTGCTTTCGATATTTTGGCCCCGTGCTGTATGTATGGTTCGAAGACTTTTATCAAACACGTAAGCGGAATCCCTGACTACTTCAAGGAGAGCTTTCCCGAAGGCTTTACTTGGGAACGTACGCAGATTTTTGAAGACGGAGGAGTTTTGACTGCCCATCAAGATACATCACTTGAGGGAAACTGTTTAATCTACAAAGTTAAGGTGTTAGGGACAAATTTTCCTGCCAATGGTCCGGTTATGCAGAAAAAGACAGCAGGCTGGGAACCGTGTGTAGAGATGTTGTACCCCCGCGACGGGGTTCTTTGCGGGCAATCCCTTATGGCCTTGAAATGTACTGATGGAAATCACTTGACATCCCATTTACGCACCACTTATCGCTCGCGTAAGCCAAGTAATGCGGTGAACATGCCGGAATTTCACTTCGGAGACCATCGTATTGAAATCTTGAAAGCCGAACAAGGAAAATTTTATGAACAATACGAATCCGCAGTGGCACGTTATTCAGATGTTCCAGAAAAGGCGACTTGATAAggcatgcaagcttggctgtt

### Sequence 6 - J23119 promoter with synthetic ribosome binding site

TTGACAGCTAGCTCAGTCCTAGGTATAATGCTAGCTGAGAAAAGAAAGGGAAACTAAGGAGGTATTTT

**SUPPLEMENTAL TABLE 1.**
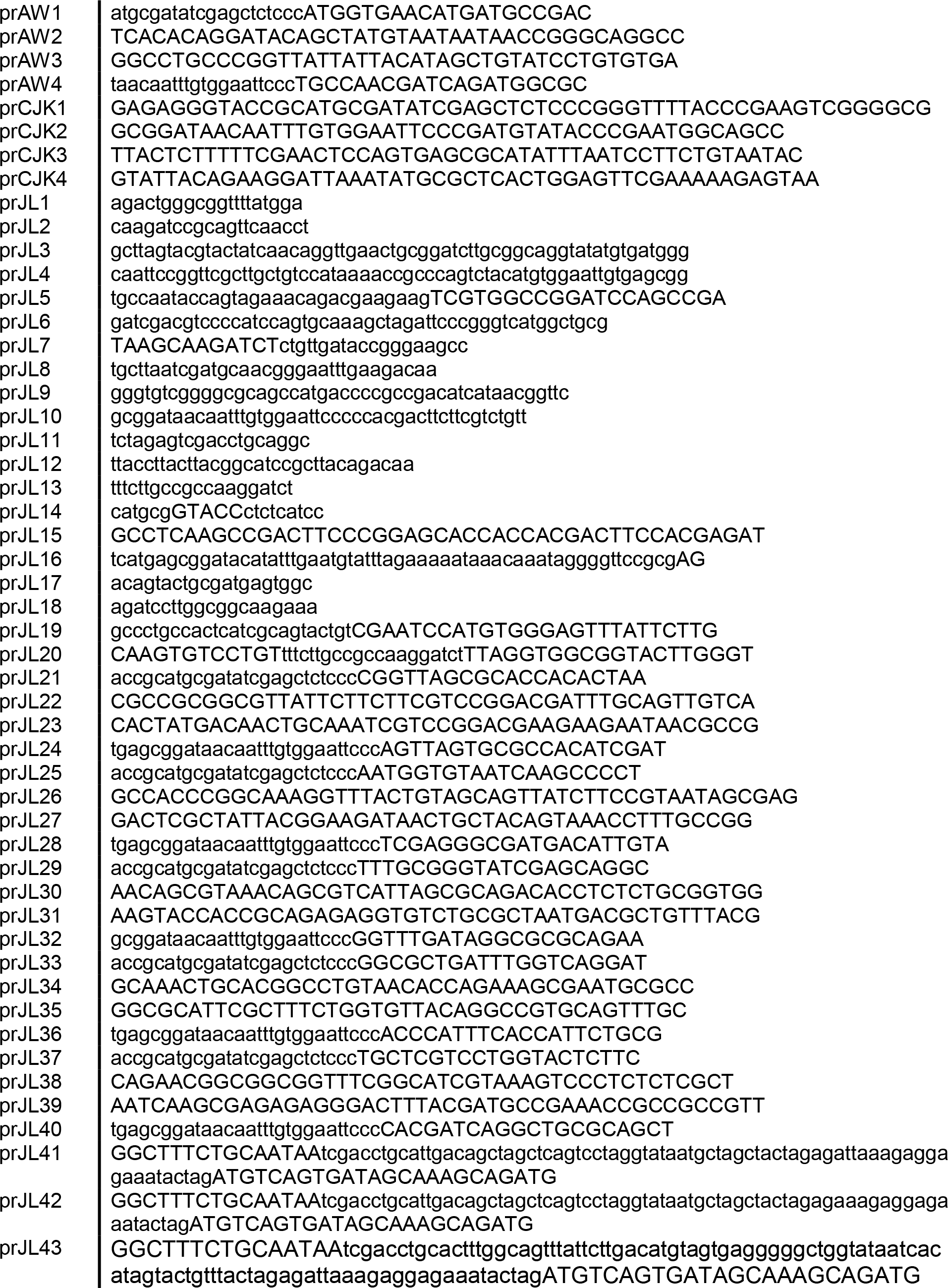
Primers used in this study

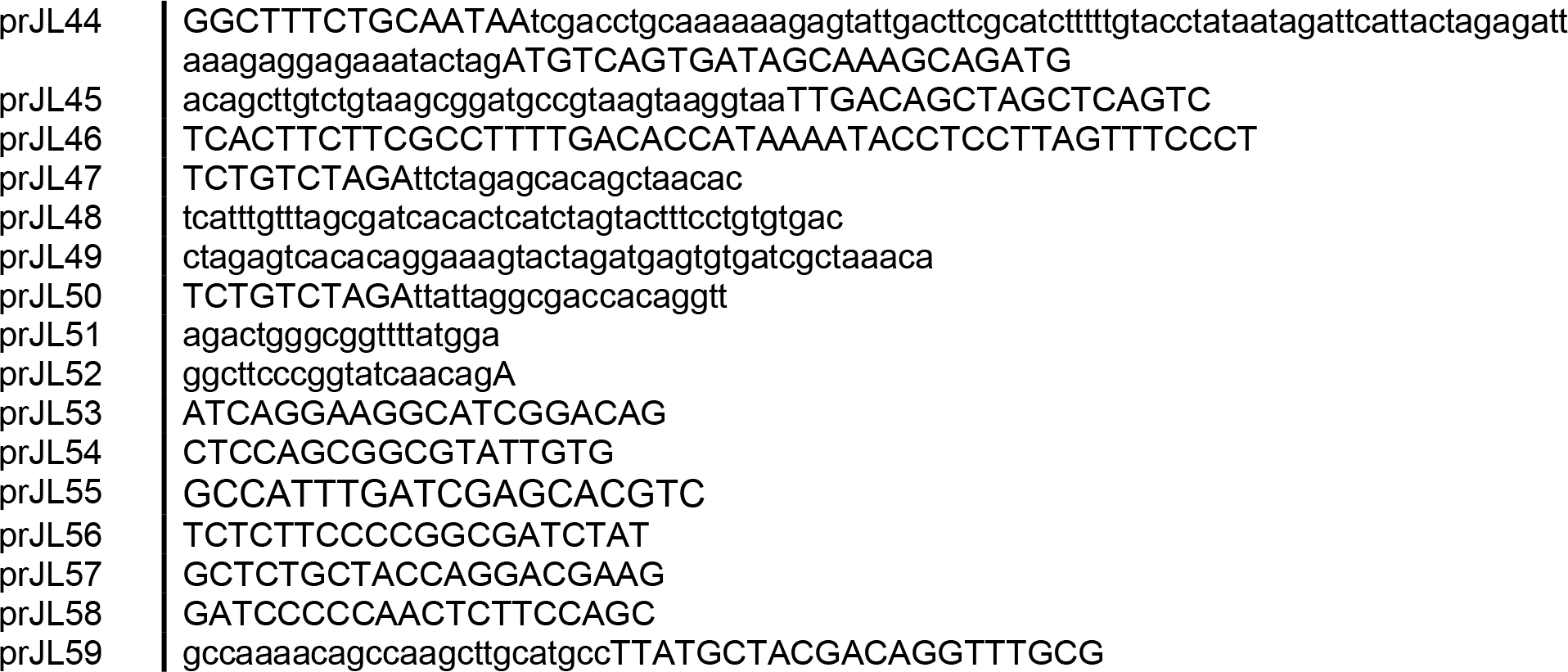

**SUPPLEMENTAL TABLE 2.**
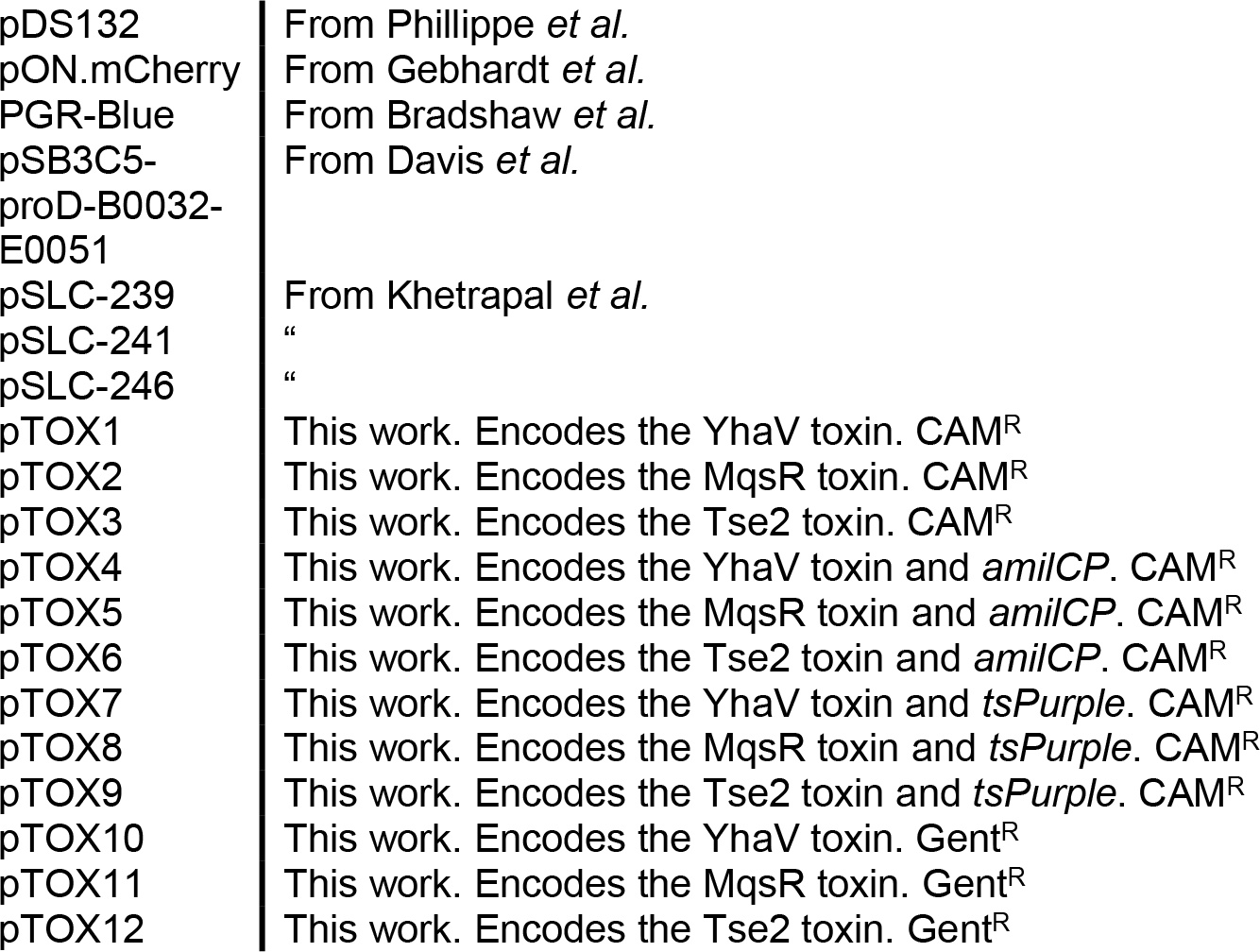
Plasmids used in this study

